# Comparative analysis of spike-sorters in large-scale brainstem recordings

**DOI:** 10.1101/2024.11.11.623089

**Authors:** Caitlynn C De Preter, Elizabeth M Leimer, Alex Sonneborn, Mary M Heinricher

**Author notes:** **Corresponding author:** Mary M Heinricher, Oregon Health & Science University, 3181 SW Sam Jackson Park Rd, Portland, OR 97239. **Author Contributions:** CCDP designed research, performed research, analyzed data, and wrote the paper. EML, AS, and MH analyzed data and wrote the paper.

## Abstract

Recent technological advancements in high-density multi-channel electrodes have made it possible to record large numbers of neurons from previously inaccessible regions. While the performance of automated spike-sorters has been assessed in recordings from cortex, dentate gyrus, and thalamus, the most effective and efficient approach for spike-sorting can depend on the target region due to differing morphological and physiological characteristics. We therefore assessed the performance of five commonly used sorting packages, Kilosort3, MountainSort5, Tridesclous, SpyKING CIRCUS, and IronClust, in recordings from the rostral ventromedial medulla, a region that has been characterized using single-electrode recordings but that is essentially unexplored at the high-density network level. As demonstrated in other brain regions, each sorter produced unique results. Manual curation preferentially eliminated units detected by only one sorter. Kilosort3 and IronClust required the least curation while maintaining the largest number of units, whereas SpyKING CIRCUS and MountainSort5 required substantial curation. Tridesclous consistently identified the smallest number of units. Nonetheless, all sorters successfully identified classically defined RVM physiological cell types. These findings suggest that while the level of manual curation needed may vary across sorters, each can extract meaningful data from this deep brainstem site.

**Significance Statement:** High-density multichannel recording probes that can access deep brainstem structures have only recently become commercially available, but the performance of open-source spike-sorting packages applied to recordings from these regions has not yet been evaluated. The present findings demonstrate that Kilosort3, MountainSort5, Tridesclous, SpyKING CIRCUS, and IronClust can all be reasonably used to identify units in a deep brainstem structure, the rostral ventromedial medulla (RVM). However, manual curation of the output was essential for all sorters. Importantly, all sorters identified the known, physiologically defined RVM cell classes, confirming their utility for deep brainstem recordings. Our findings provide suggestions for processing parameters to use for brainstem recordings and highlight considerations when using high-density silicon probes in the brainstem.

## Introduction

“Spike-sorting” refers to the process of assigning extracellularly recorded action potential waveforms, or “spikes” to distinct individual neurons. Historically, extracellular recordings have been performed using a single electrode, recording a small number of neurons, followed by semi-automated sorting based on template matching and waveform features (shape, amplitude, or width) and extensive manual curation on an individual spike basis (Gerstein and Clark 1964; Rey et al. 2015). However, the advent of multichannel recording technologies has increased data output by several orders of magnitude, making this method of sorting increasingly infeasible (Stevenson and Kording 2011; Rey et al. 2015). More fully automated spike-sorting approaches have consequently been introduced, with the goal of reducing the time, effort, and human subjectivity associated with earlier sorting techniques (Lefebvre et al. 2016). Newer sorters employ a combination of template matching, density-based approaches, and clustering, with manual curation verifying the resulting clusters (Lefebvre et al. 2016; Hennig et al. 2019; Buccino et al. 2022).

The most accurate and efficient approach for sorting a given dataset likely depends on the morphological and physiological properties of the brain region of interest. For example, recordings from brain regions with densely-packed cells with high firing rates suffer from overlapping spikes that can be assigned incorrectly during unit identification (Averbeck et al. 2006). Sorters that rely on density-based approaches have been shown to fail at resolving overlapping spikes at a higher rate than those using template-matching (Pillow et al. 2013; Garcia et al. 2022). Conversely, low firing rates can impact the performance of template-based sorters, which rely on an average waveform shape to distinguish units (Shoham et al. 2006; Pedreira et al. 2012). Therefore, the specific neuron populations in a region and corresponding firing rate distributions must be considered when choosing a spike-sorting package.

While the performance of a number of automated sorters has been evaluated and compared in recordings from the cortex, hippocampus, dentate gyrus, and thalamus (Buccino et al. 2020; Magland et al. 2020), the defined morphological cell types and layered structure in these regions gives neurons distinct electrical properties that result in distinguishable waveforms (Trainito et al. 2019). In contrast, brainstem regions, which have only recently begun to be explored at the high-density network level, have received less attention, partly due to technological challenges. Multielectrode arrays are too large to be inserted into deep brainstem structures without serious injury, and high-density silicon probes long enough to reach deep structures have only recently become commercially available (e.g. (Ulyanova et al. 2019; Shoup et al. 2024)). To date, few multichannel recordings have been reported from this region (e.g., Tsunematsu et al. 2020; Concha-Miranda et al. 2022; Malfatti et al. 2022; Strickland and McDannald 2022; Yang et al. 2023). It is therefore important to systemically assess the performance of different automated sorters in the brainstem to help identify the most effective strategies for sorting.

Given that there are differences in neuronal size, density, and firing patterns across different brain regions (Mochizuki et al. 2016), and that these might impact sorter performance, the present study compared the performance of different sorters applied to recordings from a deep brainstem region, the rostral ventromedial medulla (RVM). The RVM is a ventral brainstem region, encompassing the ventromedial aspects of gigantocellular and magnocellular reticular formation and medullary raphe, that has been well characterized using single-electrode approaches (Fields et al. 1983; Heinricher et al. 1987; Heinricher et al. 1989; Clarke et al. 1994). The different cell classes lack distinct morphology (Winkler et al. 2006), but are defined by firing changes associated with noxious-evoked withdrawal behaviors: “ON”-cells exhibit a burst of activity and “OFF”-cells a pause in activity associated with behavioral withdrawal from the stimulus (De Preter and Heinricher 2024). The third class of cells, “NEUTRAL”-cells, do not exhibit any change in activity in response to noxious stimuli. Over the last 30 years, RVM spike waveforms have been sorted using software template matching, cluster analysis, and manual verification on an individual spike-to-spike basis (Hryciw et al. 2021; De Preter and Heinricher 2023), a time- and labor-intensive approach that would be impossible in multi-channel recordings.

Here we took advantage the novel application of silicon-probe technology in RVM and the well-defined firing patterns to assess performance of these different sorters. We used SpikeInterface, a Python toolkit that integrates multiple sorters (Buccino et al. 2020), to compare performance of five different sorters, with and without manual curation.

## Methods

All animal procedures were performed in accordance with Oregon Health & Science University’s animal care committee’s regulations and followed the guidelines of the National Institutes of Health and the Committee for Research and Ethical Issues of the International Association for the Study of Pain. Male and female Sprague Dawley rats were housed in a 12-hour light-dark cycle environment with free access to water and food for at least one week prior to experiments.

### Electrophysiological recordings

Rats were briefly anesthetized (4-5% isoflurane) for external jugular vein catheter implantation. Animals were then transferred to a stereotactic frame and anesthetic plane was maintained with continuous methohexital infusion. A small craniotomy was made to gain access to the RVM and dura was removed. Following preparatory surgery, the anesthetic plane was set to maintain a stable heat-evoked paw withdrawal threshold. Heart rate and body temperature were monitored and maintained throughout the experiment. Testing was performed in low ambient light conditions (< 5 lux).

A 64-channel, high-density silicon probe was used to record RVM neuronal activity (Cambridge Neurotech M1, Cambridge, UK). Prior to placement, the probe was painted with DiI to identify probe location (Sigma-Aldrich: Cat. #42364). The probe was lowered at a rate of 1.25 micron/s using a hydraulic microdrive (David Kopf Instruments, Tujunga, CA) until the entire length (632 µm) of the contact distribution was within the RVM.

Probes were paired with a RHD 64-channel recording headstage (Intan Technologies, Los Angeles, CA) using an adaptor (ADPT A64-Om32×2, Cambridge Neurotech), and connected to both the Intan Recording Systems (RHD 1024-channel) and, in parallel, to a CED Spike2 (Cambridge Electronic Design, Cambridge, UK) data acquisition system. Signals were band-pass filtered (500 Hz to 15 kHz), sampled at 30 kHz, and stored for offline analysis.

A 25-min recording from each of six animals was used in this study. Noxious stimulation was delivered at 5-min intervals: three heat stimulations followed by a hindpaw pinch with toothed forceps. Noxious heat stimuli were applied to the plantar surface of the hindpaw using a custom-built Peltier device. The surface temperature was increased at a rate of 1.5 °C/s from 35 °C to a maximum of 53 °C. Withdrawal was determined from hamstring rectified and smoothed (0.05 s) electromyographic (EMG). EKG and core temperature were also collected.

### Histology

At the conclusion of the experiment, rats were deeply anesthetized using methohexital before being perfused intracardially with 0.9% saline followed by 4% formalin. Brains were extracted and fixed in a 4% formalin solution for 24 hours, then stored in 30% sucrose. Brains were sectioned (60 µm), and probe placement confirmed by location of DiI tracks using a fluorescence microscope (BZ-X710, Keyence Corporation of America, Itasca, IL) and plotted according to the Paxinos & Watson rat brain atlas (Paxinos and Watson 2009). Only recordings in which the entire length of the contacts (632 µm) were in the RVM were used.

### Spike sorters

We compared the performance of five established sorters on the RVM recordings: MountainSort5 (MS5) (Chung et al. 2017), IronClust (IC) (Jun et al. 2017), Kilosort3 (KS3) (Pachitariu et al. 2023), Tridesclous (TDC) (Garcia and Pouzat 2015), and SpyKING CIRCUS (SC) (Yger et al. 2018). KS3 assigns units as “good” or “mua” (multi-unit activity), and only the units labeled “good” were considered in further analyses. MS5 and IC employ a clustering algorithm, KS3 and TDC template matching, and SC a combination of clustering and template matching. Each of these sorters has been validated against “ground-truth” datasets (Buccino et al. 2020; Magland et al. 2020). Outputs from each sorter were loaded into SpikeInterface for post-processing and comparison.

### Post-processing of sorter output and comparison

The raw output of each sorter (1241 units) was post-processed (SpikeInterface postprocessing module) to eliminate units unlikely to correspond to a valid neuronal signal based on low signal-to-noise ratio (< 4.0), a high (> 0.5) interspike interval violations ratio (Vincent and Economo 2024), or few spikes (< 500). This resulted in a reduction in the of total number of unique units found by the five sorters to 671 that were used for all analyses. The post-processed output of each sorter was also manually curated in Phy (Rossant and Harris 2013). Sorted units were accepted, rejected, and split or merged to form new units (Rossant and Harris 2013; Buccino et al. 2020). Units were rejected if they were not present throughout the recording (e.g. drifted in or out during the recording), if they had contamination (e.g. two units colliding), or if they were a duplicate (e.g. units recorded from the same contacts with similar waveforms and a zero-lag cross-correlogram peak). For duplicates, only the unit with the greater number of spikes was accepted for further analysis. The curated output was then reloaded into SpikeInterface for analysis of the impact of curation.

Spike trains were compared using the SpikeComparison package of SpikeInterface. A 50% spike train match was used to extract matched units (Buccino et al., 2020). Sorter performance was compared using a Chi-square test, *t*-test, or ANOVA with Holm-Sidak *post-hoc* tests in GraphPad Prism.

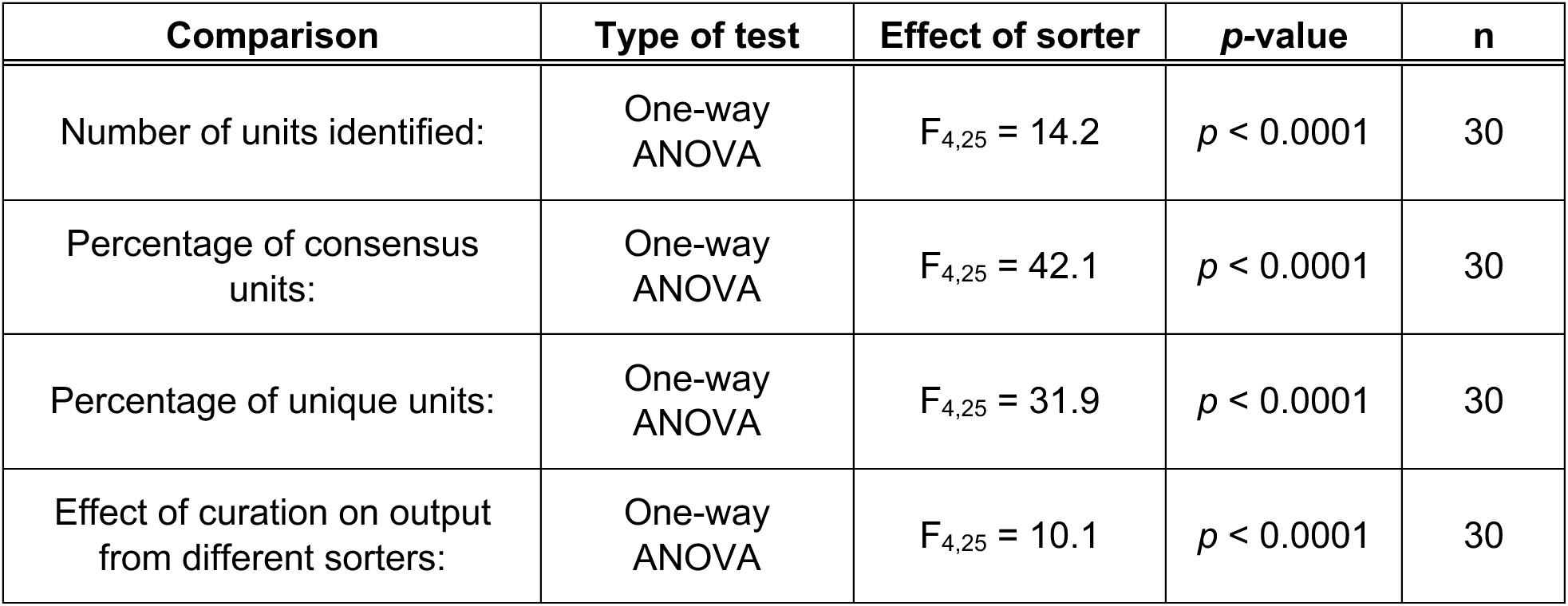
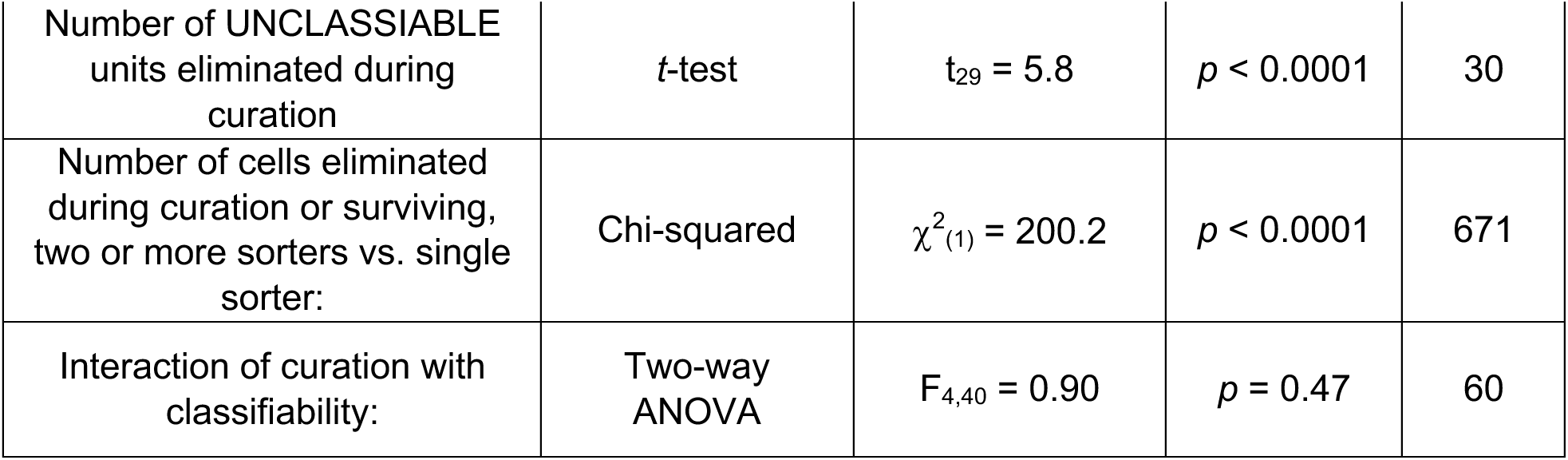

### RVM neuron functional classification

Units were classified as ON-, OFF-, or NEUTRAL-like based on change in firing rate in the 5-s interval immediately before and after onset of noxious-evoked withdrawal (Fields et al. 1983). A unit was classified as OFF-like if it exhibited an average percent *decrease* in firing rate greater than 40%, and ON-like if it showed an average firing rate *increase* greater than 100%. For units without ongoing activity, those exhibiting an increase of at least 5 spikes in the 5 s after EMG onset were also classified as ON-cells. NEUTRAL-like units had a minimum of 0.1 spikes/s and displayed no average change in firing rate greater than 50% overall, and no single trial with a decrease greater than 40% or increase greater than 100%. Units that did not match these criteria and inconsistently responded across trials were considered UNCLASSIFIABLE units.

## Results

### Comparison of five sorters

To assess the agreement between the outputs of the five tested sorters, we compared performance on six RVM recordings, from 3 male and 3 female rats. An example of units identified on 18 probe channels before and after delivery of noxious pinch to the hindpaw is shown in Figure 1A. Units had discriminable waveforms (Figure 1A, inserts) and the recording location in RVM was confirmed (Figure 1B). Of 117 units identified by at least one sorter in this recording, different sorters identified different numbers of units. SC identified the greatest number of units (70) and TDC the fewest (24). MS5, KS3, and IC identified intermediate numbers of units, with 47, 45, and 38 respectively (Figure 1C). There was also substantial variation in the degree of agreement across sorters. Of 117 total units detected by at least one sorter in this recording, 15 were identified by all five, 13% of the total (Figure 1C, red). However, these consensus units represented different proportions of the number identified by the different sorters. That is, these 15 represented almost 63% of the total identified by TDC, 39% of those found by IC, about a third of those identified by MS5 and KS3, and only 21% of those found by SC. However, another 22 units were agreed upon by two to four sorters (19% of total cells identified, Figure 1C, orange). Conversely, each sorter also identified unique units only found by that sorter (Figure 1C, yellow). TDC, which identified the fewest units overall, also identified the fewest unique units (2). IC and KS3 yielded a similar number of units not found by other sorters (7 and 11, respectively), and MS5 identified 21 unique units. SC identified 39 units that were not found by any other sorter, consistent with the large number of units identified by this sorter relative to the others. Of the 117 units identified, 80 (68%) were reported by only a single sorter, and almost half of those 80 were reported by SC.

**Figure 1.**
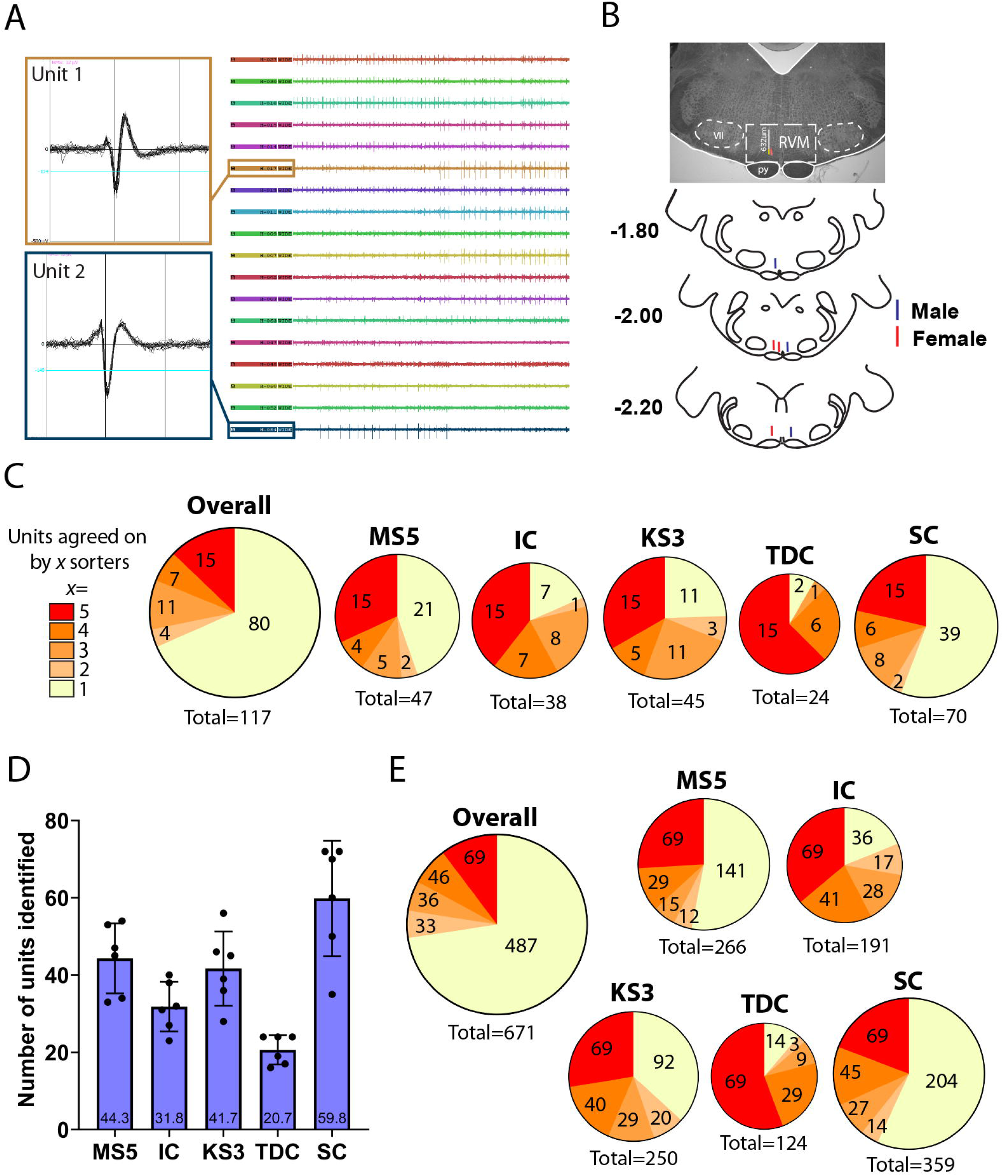
Performance of different automated sorters in brainstem recording. (**A**) Example recording. 3-s sample of spiking activity seen on 18 channels. Two example waveforms in insets. (**B**) Location of the probe. The probe was confirmed to be in RVM (632 µm, probe tip was coated with DiI (red) for visualization). py: pyramid, VII: facial nucleus. (**C**) Number of units identified by each individual sorter and across all five sorters for the example recording. Of 117 units identified by at least one sorter, 15 were agreed upon by all five, whereas 80 were found by only a single sorter. Number of sorters that agreed upon a given unit ranged from all five (red, x = 5), to only a single sorter (yellow, x = 1). Pie charts are scaled to the total number of units identified by each sorter. (**D**) Mean (± SD) number of units identified by each sorter across all 6 recordings. (**E**) Number of units identified by each individual sorter and across all five sorters summed over the six recordings. Of 671 units identified by at least one sorter, 69 were agreed upon by all five (red), whereas 487 were found by only a single sorter (yellow).

Comparison of sorter outputs across all six recordings showed that these trends seen in the example recording were consistent (Figure 1D). SC reported significantly more units than any of the other four sorters, whereas TDC identified fewer than any of the other sorters except IC (F_4,25_ = 14.2, *p* < 0.0001, n = 30). MS, KS, and IC identified intermediate numbers of units.

Of the 671 total units across all recordings that were detected by at least one sorter, 69 (10%) were agreed upon by all five sorters (Figure 1E, red, 9 to 15 units per recording). As with the example recording, these consensus units represented different proportions of the number identified by the different sorters. That is, these 69 represented over half of the total identified by TDC (57%), 36.4% of those found by IC and, 26% of identified by MS5 and 28.6% of those found by KS3, but only 20% of those found by SC. The percentage of all units identified by TDC that were consensus units was significantly greater than that for any of the other sorters, while the percentage that were consensus units was significantly less for SC than for any of the other sorters (F_4,25_ = 42.1, *p* < 0.0001, n = 30, Holm-Sidak *post-hoc* test). Another 115 (17%) were agreed upon by two to four sorters (Figure 1E, orange). By contrast, 487 (73%) were identified by only one sorter (Figure 1E, yellow). The percentage of unique units was different for the five sorters, and paralleled the total number of units identified (F_4,25_ = 31.9, *p* < 0.0001, n = 30, Holm-Sidak *post-hoc* test). That is, over half of the units identified by SC were found only by SC, whereas only about 10% of the units identified by TDC were unique to TDC.

### Effect of manual curation

A stated goal of most automated sorters is to reduce the need for manual curation. Therefore, the automated output was compared to curated output to determine which sorter likely yielded the greatest number of true units. During curation, a unit was accepted or rejected based on whether it was present throughout the recording, whether it was contaminated by a second waveform, or whether it was a duplicate unit. An example of a duplicate unit identified during curation is shown in Figure 2A. Units 21 and 22 in this example recording demonstrated similar waveform shapes and a zero-lag peak on the cross-correlogram. Unit 21 had fewer spikes and was consequently rejected as a duplicate of Unit 22.

**Figure 2.**
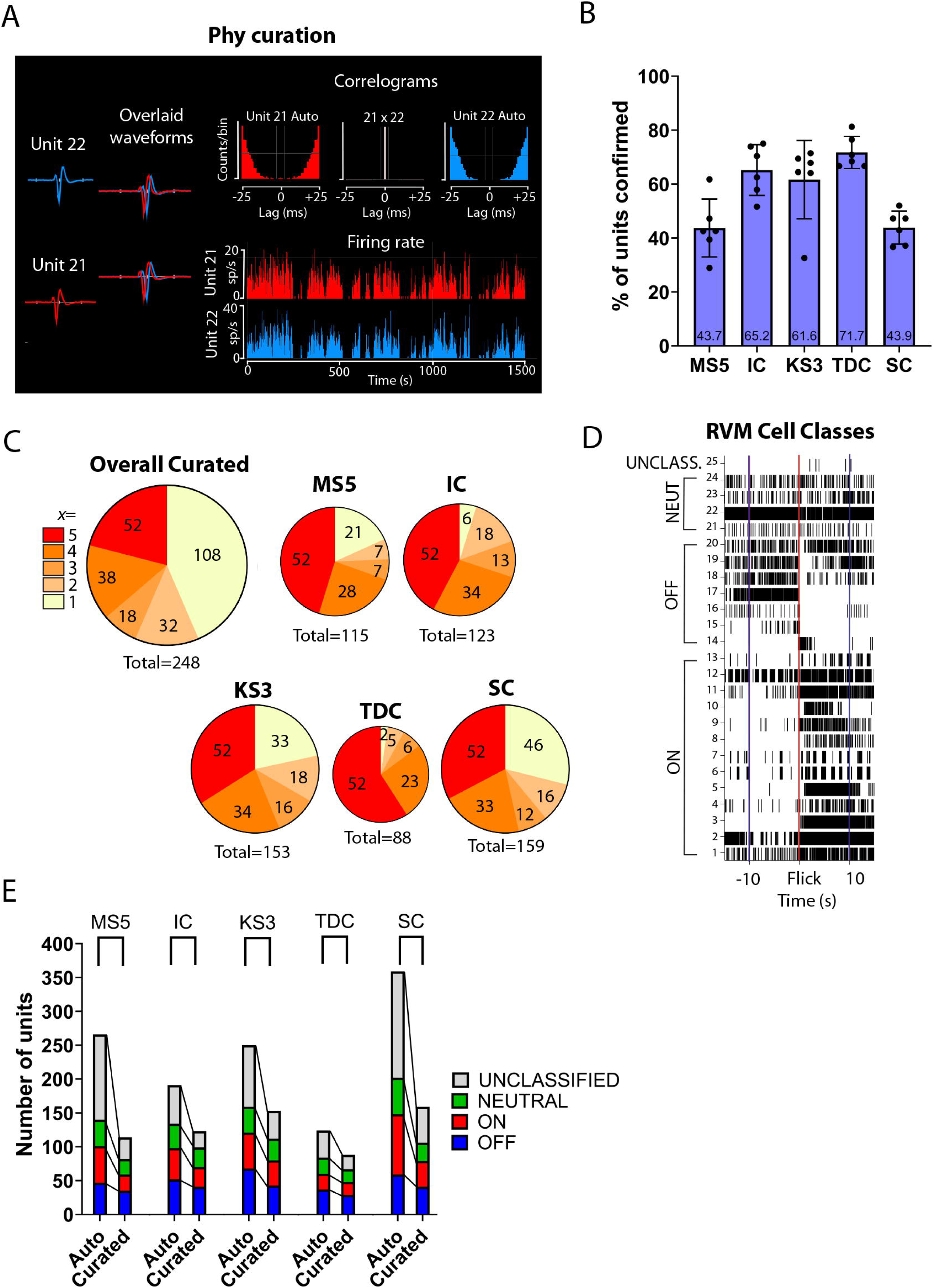
Effect of curation and interaction with physiological classification. (**A**) Example of curation of duplicate units. Unit 21 and 22 are identified as duplicates based not only on the overlapping waveform shape but on zero-lag peak in the cross-correlogram (top row, middle). Autocorrelograms (top row, left and right) show expected absence of coincident spikes. (B) Percentage of units (mean ± SD) identified by each sorter that survived curation. (C) Number of units identified by each individual sorter and across all five sorters that survived curation. Number of units agreed upon by all five sorters (red), by 4, 3, or 2 sorters (orange), or unique to a single sorter (yellow). (**D**) Example of classification of individual neurons as UNCLASSIFIED, NEUTRAL-, OFF- and ON-like. Rasterplot shows activity for 25 units identified in the curated output of KS3 during the 10 seconds before and after noxious evoked withdrawal (Flick, red line). (**E**) All sorters were able to identify neurons in the three classically defined RVM classes. UNCLASSIFIED units were disproportionately eliminated during curation. MS5 and SC identified the greatest number of UNCLASSIFIABLE units.

Of the 671 units identified in the automated output from the five sorters, 248 (37%) survived curation. Comparison of the effect of curation on the output from the different sorters showed substantial variability (Figure 2B, F_4,25_ = 10.1, *p* < 0.0001, n = 30). Thus, while TDC initially reported the smallest number of units, almost 72% of these were accepted during curation. By contrast, less than half of the units identified by MS5 and SC were accepted as valid units during curation. Considering only the 69 units originally agreed upon by all five sorters in the automated output, 52 (75%) survived curation (Figure 2C, Overall Curated, red). Of 184 units identified by at least two sorters, 136 survived curation (74%). By comparison, of the 487 unique units reported in the automated output, only 108 (22%) survived curation (Figure 2C, Overall Curated, yellow). Thus, units uniquely identified by a single sorter are less likely to survive curation that those identified by two or more sorters (χ(1) = 200.2, *p* < 0.0001). SC and KS3 identified the greatest total number of units that remained after curation, with 159 and 153, respectively (Figure 2C). IC and MS5 identified a similar number of units after curation, 123 and 115, respectively, and TDC identified 88 total units after curation (Figure 2C).

### All five sorters identify physiologically classifiable units

We next determined the ability of each sorter to identify RVM units that could be classified as ON-, OFF-, or NEUTRAL-like units. Units that exhibited changes in activity associated with noxious-evoked withdrawal can be seen in the example trials shown in raster plots (Figure 2D) before and after curation. All sorters identified both UNCLASSIFIED and classifiable RVM units (Figure 2E). Between 54% and 70% of the cells identified in the automated output were classifiable, and assigned to the ON-, OFF-, OR NEUTRAL-like classes. In the curated output, between 75% and 80% of the cells were classifiable. There was no difference amongst sorters in the percentage of classifiable units identified in the automated or curated output (two-way ANOVA, *p* > 0.05).

Although all sorters identified classifiable units, curation differentially eliminated UNCLASSIFIABLE units. As shown in Figure 2E, the numbers of both classifiable and unclassifiable units were reduced by curation. SC identified the greatest number of classifiable RVM units, with 202 total ON-, OFF-, and NEUTRAL-like units. However, curation reduced this number by almost half, to 106. The number of UNCLASSIFIABLE units was reduced by about 66%, from 157 units to 53. KS3 identified the next highest number of classifiable units with a total of 159 ON-, OFF-, NEUTRAL-like units in the automated output. Curation reduced this number by 30%, resulting in a total number of 112 units, 6 more units than SC. The number of UNCLASSIFIABLE units was reduced by about 55%, from 91 to 41. IC and MS5 reported similar numbers of classifiable units, 134 and 140 units, respectively. However, MS5 identified a much greater number of UNCLASSIFIABLE units, with 126 compared to the 57 UNCLASSIFIABLE units found by IC. After curation, the number of MS5 classifiable units was reduced by about 41% and UNCLASSIFIABLE units by around 75%, while for IC, curation resulted in a reduction of about 26% for classifiable units and 58% for UNCLASSIFIABLE units. TDC was the least impacted by curation compared to the other sorters, although it identified only 84 classifiable units prior to curation. This was reduced to 67 units after curation. The number of UNCLASSIFIABLE units was reduced by about 48%, from 40 to 21 units.

On average across sorters, there was about a 64% reduction in UNCLASSIFIABLE units but only about a 35% reduction in classifiable units following curation. Thus, across all sorters and all six recordings, curation substantially reduced the number of UNCLASSIFIABLE units, with a much smaller impact on classifiable units (*t*_29_ = 5.8, *p* < 0.0001, n = 30). In sum, all five sorters successfully identified RVM units that exhibit changes in firing that have been defined using single-electrode approaches.

## Discussion

The advent of high-density, multi-channel recording technologies has enabled the study of network level activity across brain regions. However, these advances also bring challenges for traditional spike-sorting approaches, as the increased data volume and signal complexity require new spike-sorting methods to most accurately identify individual units. The performance of different open-source sorters has been systematically evaluated and compared in recordings from cortex, hippocampus, dentate gyrus, and thalamus (Buccino et al. 2020; Magland et al. 2020). However, the relative performance of various sorters may differ in other brain regions, given that performance can be influenced by both firing patterns and the anatomical properties of the target brain region, including cell morphology, density, and arrangement of neurons (Shoham et al. 2006; Pedreira et al. 2012; Mochizuki et al. 2016; Garcia et al. 2022). Therefore, the current study addressed this knowledge gap by evaluating the performance of five open-source sorters in recordings from the rostral ventromedial medulla (RVM), a pain-modulating brainstem structure with well-characterized physiological cell classes and multiple decades of single-unit definition. Using the SpikeInterface framework, Kilosort3 (KS3), MountainSort5 (MS5), Tridesclous (TDC), IronClust (IC), and SpyKING CIRCUS (SC) were each applied to RVM recordings. Although prior studies have applied both KS3 and SC to brainstem recordings (Tsunematsu et al. 2020; Concha-Miranda et al. 2022; Malfatti et al. 2022; Strickland and McDannald 2022; Yang et al. 2023), the current study took advantage of the well-characterized physiology of RVM neurons and used the SpikeInterface framework to compare the performance of five different sorters, MS5, IC, KS3, SC, and TDC, in brainstem recordings.

### Agreement among output of different sorters applied to RVM recordings

Sorters varied widely in the total number of units identified. SC, which uses a combination of clustering and template matching (Yger et al. 2018), identified the most units, whereas TDC, which relies mostly on template matching with minimal clustering (Garcia and Pouzat 2015), consistently identified the smallest number of units. IC and MS5, which employ a clustering approach (Chung et al. 2017; Jun et al. 2017), and KS3, which uses template learning (Pachitariu et al. 2023), yielded similar numbers of units.

The five sorters also identified variable numbers of *unique* units – units not identified by any other sorter. SC not only identified the largest number of units, it also identified the largest number of unique units. Although IC, KS3, and MS5 yielded similar numbers of units overall, MS5 found more unique units.

Performance of sorters might be influenced by anatomical and physiological differences that contribute to either too few spikes to resolve a unit, which impacts template-based sorters, or overlapping spikes, which impacts density-based clustering sorters. The medial reticular core differs significantly from cortical and hippocampal regions in terms of cellular organization. Unlike the layered cortical and hippocampal structures with distinct morphological cell types creating varied electrical properties that result in relatively distinguishable waveforms (Trainito et al. 2019), the RVM is marked by medium to large multipolar neurons compressed in the rostro-caudal plane, giving a “stacked poker chip” organization (Scheibel and Scheibel 1967; Humphries et al. 2006). Additionally, the RVM functional classes do not have distinct morphological features that would contribute to characteristic extracellular action potential waveforms (Winkler et al. 2006). Nonetheless, the variation in the total number of units, agreement amongst sorters, and number of unique units found by each sorter is not inconsistent with a previous analysis of sorters applied to a single recording spanning cortex, hippocampus, dentate gyrus, and thalamus (Buccino et al. 2020). Based on both manual curation of their sample recording and on analysis of a simulated dataset, for which ground-truth was available, these authors argued that units agreed upon by more than one sorter are likely real, whereas unique units are more likely false positives. In the present study, about 27% of all units identified in the automated output from the five sorters were detected by at least two of the sorters, and units agreed upon by at least two sorters were more likely to survive manual curation, suggesting these units likely correspond to real units.

One false-positive that was observed across sorters was the identification of duplicate units. Duplicate units arise when a spike is assigned to multiple clusters, due to slight shifts in waveform shape (Dehnen et al. 2021). This is problematic in densely packed regions like the brainstem, where spikes from neighboring neurons or from different parts of the same neuron (e.g. somata, dendrites) overlap frequently. The presence of duplicates in all sorter outputs highlights the necessity of careful manual curation to prevent duplicate units from artificially inflating unit counts and distorting interpretations of firing dynamics.

An additional factor that could influence the sortability of recordings from different brain regions is probe geometry, as contact spacing and layout influence the ability to resolve distinct units. Indeed, while the goal of the present study was to compare performance of different sorters applied to recordings from a brainstem site with well-characterized physiological properties, it could be useful to assess performance of these same sorters on recordings with this probe in different brain regions to determine whether and how probe geometry interacts with the sorter. This could also help determine whether certain probes geometries are more effective in deep brain structures and guide future development of recording technologies.

### All sorters identified classifiable RVM units

The mutually exclusive and exhaustive OFF/ON/NEUTRAL-cell framework for classification of RVM neurons is based on noxious event-related changes in firing, with OFF-cells exhibiting a pause in firing and ON-cells a burst associated with nocifensive withdrawal. NEUTRAL-cells are defined by exclusion, failing to show either a pause or a burst associated with nocifensive behaviors (Fields et al. 1983; Heinricher et al. 1989). Units corresponding to each of these three classes were identified by all sorters, and present in both the automated and curated output of each sorter.

Given the robust classification of RVM neurons in single-electrode recordings, and despite identification of OFF-, ON-, and NEUTRAL-like units in our multichannel recordings, it may be surprising that we also identified units that could not be classified. Units were considered UNCLASSIFIABLE either because they lacked sufficient activity to characterize possible responses or because apparent responses were inconsistent. The presence of UNCLASSIFIABLE units thus likely reflects the difficulty of fully characterizing each individual unit in a multi-channel recording. The single-electrode approach allows an investigator to optimize stimulus delivery so that changes in firing will be visible. That is, a “pause” in firing can only be seen during periods when the unit to be classified is spontaneously active, whereas a “burst” would be most evident only when the unit is not spontaneously active. The single-electrode approach allows full characterization of an individual unit, but is not feasible with a multi-channel recording, in which spontaneous firing can vary across different channels at different times. We therefore used a relatively insensitive measure, average change in firing rate, to classify an individual unit as OFF-, ON-, or NEUTRAL-like. With that approach, an OFF-cell with low ongoing activity or an ON-cell with high ongoing activity would have at best inconsistent changes in firing rate, causing it to be categorized as UNCLASSIFIABLE here. More sustained noxious stimulation or pharmacological interventions, such as morphine, which reliably activates OFF-cells and suppresses firing of ON-cells (Fields and Heinricher 1985; Hryciw et al. 2021), may be necessary to fully and accurately classify RVM neurons in high-density recordings.

Interestingly, the number of UNCLASSIFIABLE units was preferentially reduced by curation: overall, by about two-third. By contrast, the number of classified (OFF/ON/NEUTRAL-like) units was reduced by only about a third. This suggests that UNCLASSIFIABLE units more frequently represented false-positives, whereas “real” units more commonly exhibit firing patterns consistent with what has been reported with single-electrode approaches. The slight reduction in classifiable units during curation was not a limitation. Indeed, one false-positive that was observed in both classifiable and UNCLASSIFIABLE groups and across sorters was duplication, which could lead to incorrect conclusions about population coding and dynamics in this region. Duplicate units arise when a spike is assigned to multiple clusters, presumably due to slight shifts in waveform shape. If not ruled out in curation, duplicate units would artificially inflate the total unit count and distort interpretations of firing dynamics.

### MS5, IC, KS3, SC, and TDC can all be used to sort high-density RVM recordings

In the present study, MS5 required the most amount of curation, with 57% reduction in classified units, and about 75% of UNCLASSIFIABLE units eliminated during curation. SC required a similar level of curation, with more than half of all units eliminated during curation. IC, KS3, and TDC required less curation. Almost three-quarters of units identified by TDC survived curation, and this sorter also identified the smallest number of UNCLASSIFIABLE units. However, it also consistently identified the smallest number of units compared to the other sorters. IC identified the second-smallest number of UNCLASSIFIABLE units and curation resulted in a relatively small decrease in the number of classifiable units. For KS3, over a third of units were eliminated during curation. However, this sorter identified the greatest number of classifiable units that survived curation. KS3 and IC thus produced the greatest number of classifiable RVM units with less intense curation.

### Conclusions

Any method for assessing activity of a neuronal population necessarily samples a subset of that population. Extracellular recording reveals only neurons that are active or for which there is a search stimulus, and with action potentials that can be resolved with a particular electrode technology. This depends both on the properties of the electrode and of the cell population under study including packing density, morphology of individual cells, and their arrangement (Robinson 1968; Lemon 1984). Choice of sorter is thus one of many factors that will influence which cells are “seen” using a given experimental protocol. Parallel limitations apply in use of calcium imaging, where expression of the indicator, optical constraints, thresholding, and selection based on activity define the subset of the relevant population that is sampled (Papaioannou and Medini 2022). Thus, although different sorters tested here revealed different subsets of the RVM population, any of the sorters in this study could reasonably be used to sort high-density brainstem recordings, albeit with varying degrees of curation efforts.

The present study highlights some considerations that will be important in any application of multi-channel recording technologies. Investigators should explicitly report how units were accepted for further study. Further, analyses of both ongoing and evoked firing patterns will be more accurate if the experimental protocol is informed by “ground truth” understanding of the neurophysiological properties of system under study. However, focusing on those units thought to be relevant to the research question should be balanced by consideration of units that might exhibit potentially interesting, but new, firing patterns. Finally, consensus amongst sorters appears to improve confidence in results in brainstem recordings, as shown previously in forebrain (Buccino et al. 2020).

**Table 1.** Statistical analysis results for effect of sorter and manual curation on number of units for brainstem recordings.

## Notes

**Funding sources:** This study was supported by the National Institutes of Health Grant NS098660 (to M.M.H). C.C.D.P. was supported by the National Institute of Dental and Craniofacial Research Fellowship F31 DE030677.

### Competing Interest Statement

The authors have declared no competing interest.

